# Skill acquisition and gaze behavior during laparoscopic surgical simulation

**DOI:** 10.1101/2020.07.17.206763

**Authors:** Sicong Liu, Rachel Donaldson, Ashwin Subramaniam, Hannah Palmer, Cosette Champion, Morgan L. Cox, L. Gregory Appelbaum

## Abstract

**Background:** Experts consistently exhibit more efficient gaze behaviors than non-experts during motor tasks. In surgery, experts have been shown to gaze more at surgical targets than surgical tools during simple simulations and when watching surgical recordings, suggesting a proactive control strategy with greater use of feedforward visual sampling. To investigate such expert gaze behaviors in a more dynamic and complex laparoscopic surgery simulation, the current study measured and compared gaze patterns between surgeons and novices who practiced extensively with laparoscopic simulation.

**Methods:** Three surgeons were assessed in a testing visit and five novices were trained and assessed (at pre-, mid-, and post-training points) in a 5-visit protocol on the Fundamentals of Laparoscopic Surgery peg transfer task. The task was adjusted to have a fixed action sequence to allow recordings of dwell durations based on pre-defined areas of interest (AOIs). Novices’ individualized learning curves were analyzed using an inverse function model, and group-level differences were tested using analysis of variance on both behavioral performance and dwell duration measures.

**Results:** Trained novices were shown to reach more than 98% (*M* = 98.62%, *SD* = 1.06%) of their behavioral learning plateaus, leading to equivalent behavioral performance to that of surgeons. Despite this equivalence in behavioral performance, surgeons continued to show significantly shorter dwell durations at visual targets of current actions and longer dwell durations at future steps in the action sequence than trained novices (*ps* ≤ .03, Cohen’s *ds* > 2).

**Conclusion:** This study demonstrates that, whereas novices can train to match surgeons on behavioral performance, their gaze pattern is still less efficient than that of surgeons, suggesting that eye-tracking metrics might be more sensitive than behavioral performance in detecting surgical expertise. Such insight can be applied to develop training protocols so non-experts can internalize experts’ “gaze templates” to accelerate learning.

**Article Summary:** Gaze pattern differences persist between laparoscopic surgery experts and novices who have been trained to reach over 98% of individualized behavioral learning plateaus in the Fundamentals of Laparoscopic Surgery (FLS) peg transfer task.

The importance of this finding lies in motivating the decision and method of including gaze behaviors via eyetracking technology in the present surgical training programs.

---

Laparoscopic surgery is a type of minimally invasive surgery in which narrow tubes are inserted into the body through small incisions, allowing surgeons to manipulate, cut and sew tissue with relatively less trauma, leading to faster patient recovery and lower morbidity compared to open surgical techniques^1^. Although ultimately beneficial for the patient, laparoscopic surgery can create challenges for surgeons. One such challenge is that laparoscopic surgeons cannot directly view the tissue they are operating on but instead must view two-dimensional video, captured by the laparoscope inserted inside the body and projected to a display at eye level through a closed-circuit camera. In addition, laparoscopic surgeons must deal with the “fulcrum effect”, whereby the tips of the surgical tools move in the opposite direction of tool handles with little tactile feedback^2^. Therefore, successful laparoscopic surgery entails expertise in depth perception from twodimensional images, as well as complex visual-motor coordination and transformation, among other important skills. Even though all these skills can be reasonably trained in existing and validated simulation programs^3^, such training takes a substantial amount of time from surgical trainees who are regularly fatigued from other professional commitments and face restricted working hours^4^. An important need therefore exists to further improve the efficiency of training programs in laparoscopic surgery, motivating research to characterize the gaps between experts and non-experts and innovation to create effective interventions to minimize such gaps.

One line of research that has showed promise for elucidating surgical expertise is the use of eye-tracking technology^5,6^. Eye-tracking is particularly well-suited as a research tool in laparoscopic surgery as it takes full advantage of the range of attentional focus defined by the monitor that displays monocular images at approximately eye level. Past research comparing gaze patterns between experts and non-experts has revealed that laparoscopic expertise can not only be detected with behavioral metrics, such as task completion times, but also on “eye” metrics in simulation tasks. Relative to non-experts, laparoscopic surgery experts consistently gaze more at the surgical targets (i.e., elements to be manipulated) than surgical tools (i.e., instruments used to manipulate) in mostly one-handed simulation tasks^7–9^ and when watching surgical recordings^10^. Because gazing at upcoming surgical targets represents a feedforward sampling strategy, whereas gazing at currently engaged targets or surgical tools represents an online sampling strategy, such a finding suggests a proactive attentional profile among experts. This insight has been applied to developing training protocols that use either prerecorded or simultaneous gaze behaviors from experts so that non-experts can recognize and learn a “gaze template” during practice, thereby promoting proactive and feedforward behavior that may lead to faster learning and better performance. Evidence from these studies supports the feasibility and validity of this gaze-based training approach. However, tasks that have been tested thus far are limited to relatively simple surgery simulations that demand mostly single-handed movements.

Because continuous bimanual coordination is essential to the success of laparoscopic surgery, further insights can be gleaned by exploring gaze behaviors in complex and dynamic bimanual laparoscopic tasks. One such task is the peg transfer task from the Fundamentals of Laparoscopic Surgery (FLS) training program, whose criterion-based completion is required for all surgery residents by the American Board of Surgery^15^. The peg transfer task emphasizes bimanual coordination by using two Maryland Dissectors in a procedure of moving six plastic objects, in turn, from the left to the right side of a flat pegboard, and later reversing the entire process to move the objects back to the left side. To be successful at this task, one must attend to the transfer of the object between the two dissectors, then the placement of the object onto the target peg to make sure that it arrives flush on the pegboard, before looking ahead to the next object that will be moved and engaging it. This validated task simulates the critical action of transferring and positioning a needle between needle holders in suturing^3^. An additional strength of considering the peg transfer task is that behavioral learning curves on this task have been shown to be fit well by an inverse function model that is capable of estimating one’s “learning plateau”, the theoretical best score one can reach with an unlimited amount of practice^16^. Therefore, when coupled with eye-tracking, the peg transfer task offers the opportunity to explore differences in gaze patterns between experts and non-experts and understand how gaze patterns evolve in individuals as they practice and gain proficiency.

The present study attempted to accomplish these goals by examining gaze and behavioral patterns during the peg transfer task. Comparisons were made between experienced surgeons and novice participants with no surgical experience, prior to, and after, they completed a training paradigm designed to meet a theoretically derived behavioral learning plateau. For this purpose, dwell-based gaze metrics, which can be regarded as one’s perceived area of importance^5^, were extracted within a set of pre-defined areas-of-interest (AOIs) that helped quantify online and feedforward sampling during timed task performance. Based on previous research, greater feedforward sampling control was expected in laparoscopic surgery experts, relative to novices, prior to practice. As novices learned and approached their behavioral learning plateau, however, it was expected that they would show similar gaze patterns to surgeons by demonstrating greater feedforward sampling control.

## Methods

### Participants

Three experienced surgeons (2 female) and five novices (3 female) participated in this study. Sample size was determined based on previous eye-tracking and behavioral research^10,17^. All three surgeons held faculty positions in the Department of Surgery within the Duke University School of Medicine with specialties in surgical oncology and bariatric surgery, with seven, two, and eight years of post-fellowship independent surgical practice, individually. All surgeons reported prior performance with the FLS peg transfer task and reported right hand dominance in their surgical activities. Their average age was 41 (*SD* = 1.73) years. All novice participants were right-handed with an average age of 26.4 (*SD* = 3.43) and none had prior experience with laparoscopic surgery or the FLS curriculum. Informed consent was given by all participants at the start of participation, and the research protocol was reviewed and approved by the Duke Health Institutional Review Board (Pro00078782). All participants were compensated at the rate of $15/hr.

### Task and Apparatus

As illustrated in **Figure 1A,** the peg transfer task was conducted on a cart mounted FLS trainer box (Limbs & Things Ltd., Savannah, GA). In order to provide a standardized procedure for peg transfers, the objects were labeled one through six, as was the pegboard at the base of the pegs on the left and right sides of the board **(Figure 1B).** Consistent with instructions provided in the FLS curriculum tutorial video^1^, participants were instructed to move the objects, one at a time, from the numbered peg on the left side, to the corresponding numbered peg on the right side. Once all six objects were placed on the right side, they then reversed this procedure and returned the objects to the left, always in order from one to six. For each of these transfers, the objective was to be picked up with the dissector on the same side, and passed in midair to the other dissector, before placing it down on the appropriate peg. As such, one repetition of the task is divided into 12 transfers, each of which begins when the dissector touches the object at the starting peg and ends when the object is lowered on the target peg and is flat on the surface of the pegboard.

**Figure 1.**
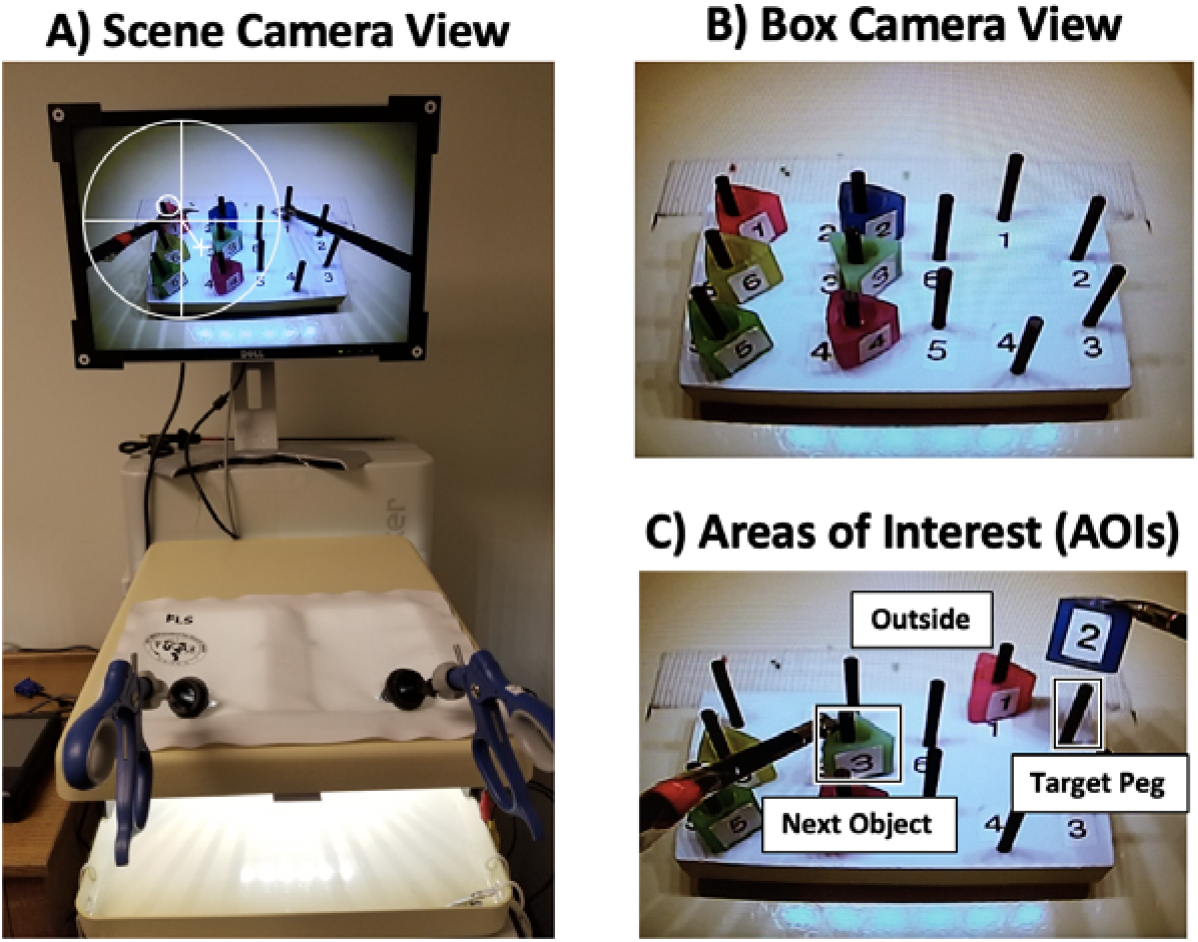
(A) Study apparatus as seen from the scene camera with pupil tracking (large circle and cross) and gaze location (small circle) indicated, neither of which are visible to the participant. (B) Pegboard as seen from the FLS camera view. (C) Example areas of interest (AOIs) during the 2^nd^ transfer of the task.

The Argus ETMobile system (Argus Science, Tyngsborough, MA) was used to track foveal vision at 30 Hz. The eye-tracker featured eye and scene cameras mounted on a pair of light-weight glasses that compute gaze locations using both camera recordings via the pupil-to-corneal-reflection technique. This technique relies on modeling spatial relationships between the black pupil and mirror reflections of three infrared lights from the cornea front surface. The gaze point is represented by a circular cursor spanning 1 of visual angle on the scene camera recordings. The eye-tracking system was further set up so that recordings from the closed-circuit camera installed inside the FLS trainer box was recorded to the same eye tracking software system. Post processing allowed spatial and temporal alignment of both scene and trainer box camera recordings so that gaze locations could be calculated by transferring coordinates from the scene camera recording to the trainer box recording. The gaze locations on the trainer box recording were subsequently used to calculate fixations, defined as a single gaze of at least 100 ms within 1 of visual angle.

### Measures

The primary eye-tracking metric used in this study was the *Percent Dwell Duration.* This normalized dwell measure was calculated by dividing the dwell duration in which consecutive fixations remain in a given AOI by the total dwell duration recorded in the corresponding transfer. Two square-shaped AOIs were defined **(Figure 1C)** and interpreted for each transfer including:

1. AOI_TP_ represents the region including and surrounding the *Target Peg.* Fixations here reflect 1-step, feedforward sampling control.
2. AOI_NO_ represents the region including and surrounding the *Next Object* to be transferred. Fixations here reflect 2-step, feedforward sampling control.
3. AOI^Outside^ represents the region excluding the AOI_TP_ and AOI_NO_. Fixations in this AOI are outside of the other regions and primarily correspond to gaze on the currently moving object, reflecting online sampling control during movement.

Because the 12^th^ transfer is the end of the repetition and does not have an AOI_NO_, it is not included in the calculation of scores. To test the consistency of the AOI definitions, the AOI sizes (pixel^2^) for both AOI_TP_ and AOI_NO_ were extracted and tested between the surgeon and novice groups, resulting in non-significant differences (*ps* > .39).

The behavioral *Performance Score* followed previous research^18^ and was computed by accounting for both task completion time and errors. Errors included drops within the field-of-view, drops outside of the field-of-view, and improper transfers (e.g., using the wrong dissectors to move an object or resting the object on a peg during transfer), and were penalized by increased task completion time (seconds). Specifically, drops within the field-of-view and improper transfers added one second each to the completion time, while drops outside the field-of-view added three seconds each to the completion time, reflecting a heavier penalty for this more serious error. The final task completion time was converted into a performance score, by dividing, if applicable, the penalized task completion time by 12 which is the number of objects transferred. The performance score thus adopts the unit of ‘seconds per object’ transferred with lower scores corresponding to better performance.

### Procedure

All participants underwent identical acclimation procedures at the start of the study. Specifically, they heard a brief verbal description of the procedures, gave informed consent, and completed a demographic survey, prior to watching an instructional video of the FLS peg transfer task and familiarization with the instruments. After this acclimation, the study activities differed between the two groups **(Figure 2).**

**Figure 2.**
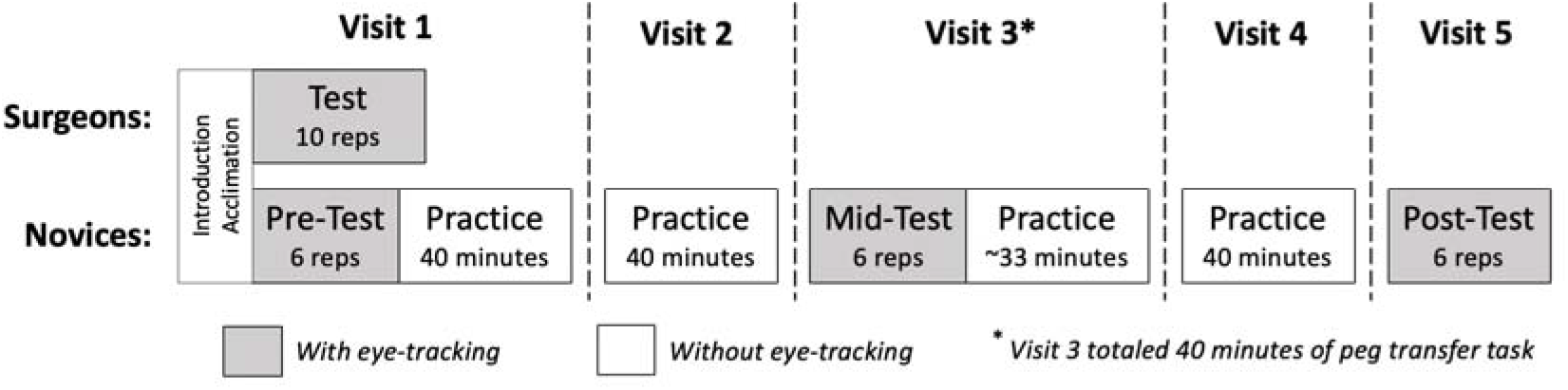
Study procedures illustraing common introduction and acclimation for both groups, as well as timed test periods with eye tracking recorded shown in gray and untimed practice periods without eye tracking shown in white.

During the remainder of their only experimental visit, surgeons were asked to complete 10 repetitions of the task as quickly and accurately as possible while wearing the eye-tracker. These repetitions were timed and constituted their performance test. Novices, however, proceeded to complete both testing (with the eyetracker) and training (without the eye-tracker) over five visits that occurred within two weeks. Following introduction and acclimation in Visit 1, novices completed a *pre-test* in which they performed six timed repetitions of the peg transfer task as quickly and accurately as possible while the eye tracker recorded their gaze. The eye tracker was then removed and novices were given a 5-minute break prior to completing two 20minute practice blocks with a 5-minute break between the blocks. During both Visit 2 and 4, novices practiced two 20-minute blocks, separated by a 5-minute break, but did not perform any testing with the eye tracking system on. Visit 3 began with a *mid-test* in which they again completed six timed repetitions as quickly and accurately as possible as eye-tracking was worn to record their gaze. This was followed by approximately 33 minutes of practice without the eye tracker to round out a total of 40 minutes of exposure to the peg transfer task on this visit. Visit 5 consisted of only a *post-test* during which novices performed six timed repetitions of the task as quickly and accurately as possible with the eye tracker to record their gaze.

### Analysis

R^19^ and JASP (vθ.11.1) were used for statistical analyses. For novices, the effect of training was evaluated using both individualized learning plateaus estimated with an inverse function model^16^ and changes across testing sessions using ANOVAs. Specifically, a 6 (Repetitions: 1 through 6) by 3 (Session: pre-test, mid-test, post-test) ANOVA was run with performance scores, and a 3 (AOI: outside, target peg, next object]) by 3 (Session) ANOVA was performed with percent dwell duration. To compare surgeons to novices, both prior to and after training, group differences on performance score were tested using 6 (Repetition) by 2 (Expertise: surgeons, novices) ANOVAs, and group differences on percent dwell duration were tested using 3 (AOI) by 2 (Expertise) ANOVA^2^. In both of these analyses, only the first six repetitions of the surgeons’ test were used in order to match the six repetitions collected with the novices during their tests, given that no statistical differences were identified across all the 10 repetitions for the surgeons. In order to further investigate gaze differences and determine if subtle differences in the quantification of dwell patterns influenced the findings, identical ANOVA analyses were also performed using an alternative dwell metric, Percent Dwell Count, whose results can be found in the **Supplement.** The Greenhouse-Geisser correction on degree-of-freedom was used when Mauchly’s test for sphericity reached statistical significance. When post-hoc pairwise comparison was needed, the familywise alpha level was controlled using the Holm-Bonferroni method. The alpha level was set at .05.

## Results

### Individually Modeled Learning Curves

The total amount of time spent performing the peg transfer task (including time spent testing) ranged between 171 and 177 (*M* = 173, *SD* = 2.71) minutes among the five novices. As shown in **Figure 3,** the inverse function model fit well to the behavioral performance data from the five novices, *ps* ≤ .02, R^2^s ≥ .30. Specifically, the results indicated that all of the novices spent less than 60 (*M* = 24.40, *SD* = 18.88) minutes performing the task to reach 90% of their behavioral performance plateaus, and all reached approximately their estimated plateau by the end of training (*M* = 98.62%, *SD* = 1.06%).

**Figure 3.**
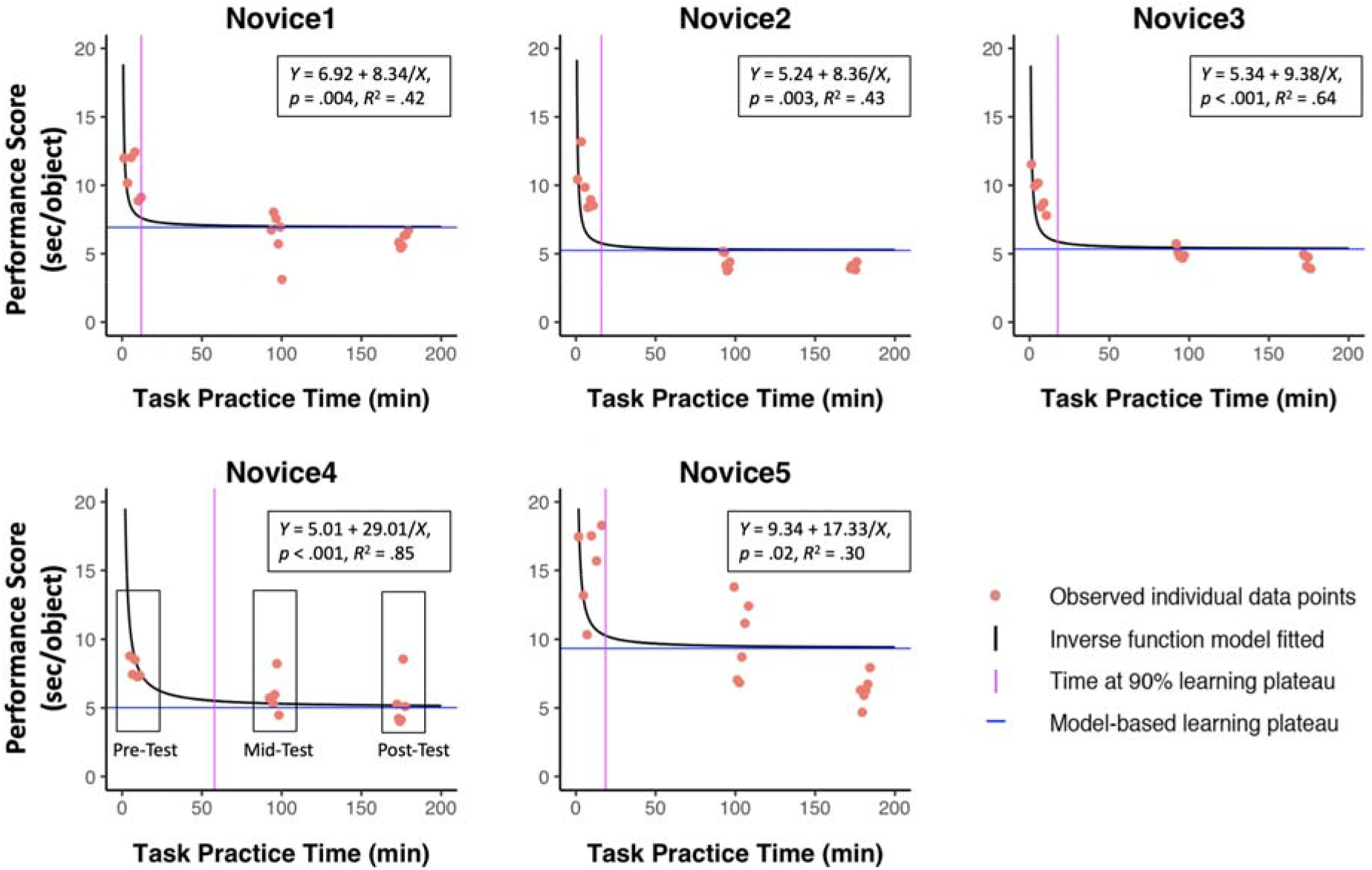
Individual data points, learning curves, inverse model functions and plateaus for novices across training

### Novice Learning across Test Sessions

Repetition by Session ANOVA performed on the seconds-per-object performance scores **(Figure 4A)** indicated a significant main effect of Session, *F*_(2,8)_ = 50.58, *p* < .001, *η*^2^_*p*_ = .93. Pairwise comparisons demonstrated that novices improved significantly from pre-test (*M* = 11.11, *SD* = 3.85) to mid-test (*M* = 6.33, *SD* = 2.51), *p* < .001, Cohen’s *d* = −7.77, and from pre-test to post-test (*M* = 5.20, *SD* = 1.86), *p* = .004, Cohen’s *d* = −3.21, but not from mid-test to post-test, *p* > .15. No significant main effect of Repetition or Repetition by Session interaction were observed (ps > .16).

For the percent dwell duration metric **(Figure 4C),** the AOI by Session ANOVA indicated a significant effect of AOI, *F*_(2,8)_ = 84.20, *p* < .001, *η*^2^_*p*_ = .96. Pairwise comparison showed significantly higher percent dwell duration for AOI_Outside_ (*M* = 73.6%, *SD* = 9.5%) than for AOI_NO_ (*M* = 13.9%, *SD* = 4.4%), *p* < .001, Cohen’s *d* = 8.06, and AOI_TP_ (*M* = 12.4%, *SD* = 5.9%), *p* < .001, Cohen’s *d* = 7.73. No significant main effect of Session or Session by AOI interaction were observed (ps > .10).

**Figure 4.**
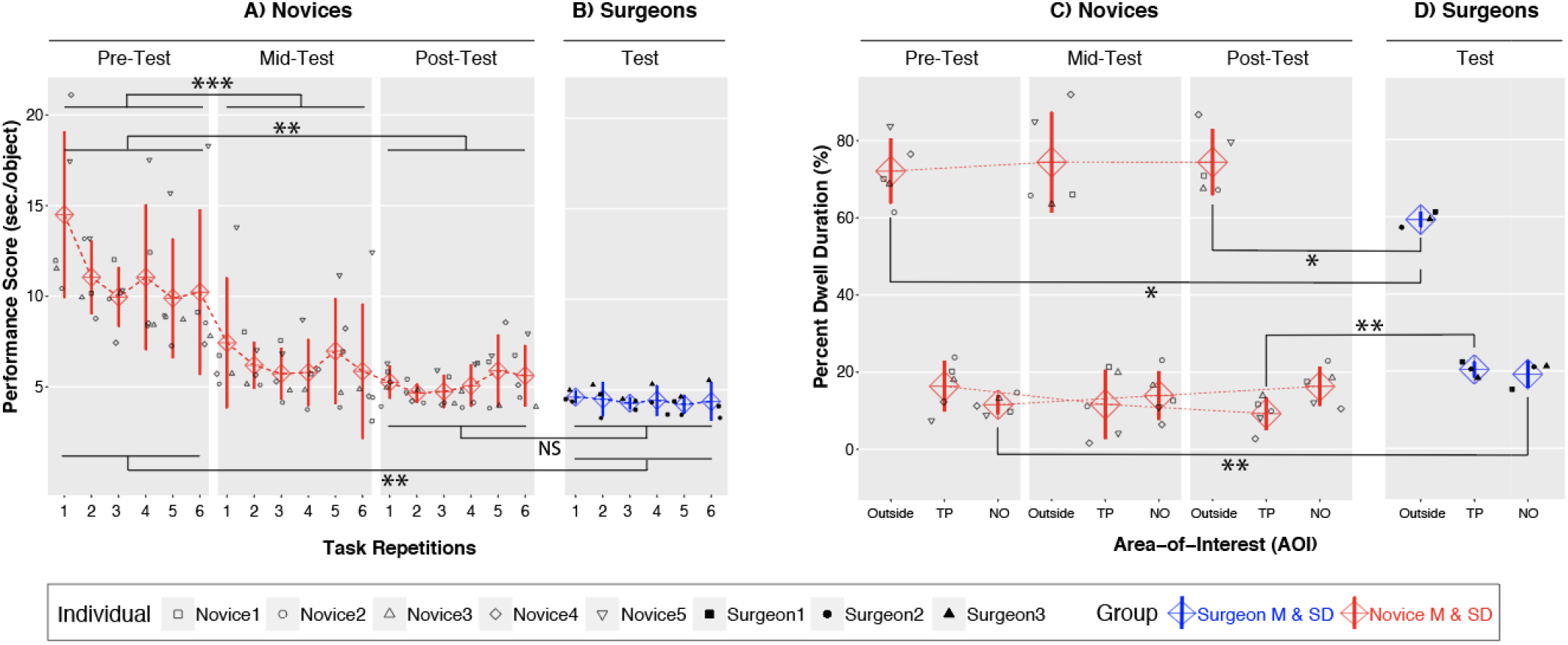
Behavioral performance and percent dwell duration (for both Individuals and groups) with important comparisons statistically marked. (A) Novice performance scores for each repetition across the pre-, mid- and post-test. (B) Surgeon performance scores for each repetition. (C) Novice percent dwell duration for each AOI shown for each testing session. (D) Surgeon percent dwell duration for each AOI. ***p < .001, **p < .01, p < .05, NS. non-significant.

### Surgeons versus Novices

When comparing performance score between novices at pre-test **(Figure 4A, left)** and surgeons **(Figure 4B),** the Expertise by Repetition ANOVA showed a significant main effect of Expertise, *F*_(1,6)_ = 20.55, *p* = .004, *η*^2^_p_ = .77, demonstrating better performance scores in surgeons (*M* = 4.39, *SD* = 0.57) than novices (*M* = 11.11, *SD* =2.45), Cohen’s *d* = −3.77. This same comparison between novices at post-test **(Figure 4A, right;** *M* = 5.20, *SD* = 0.99) and surgeons **(Figure 4B)** was not significantly different (*ps* > .24).

When comparing percent dwell duration of novices at pre-test **(Figure 4C, left)** and surgeons **(Figure 4D),** the Expertise by AOI ANOVA revealed a significant main effect of AOI, *F*_(2,12)_ = 138.71, *p* < .001, *η*^2^_p_ = .96, and a significant Expertise by AOI interaction, *F*_(2,12)_ = 5.20, *p* =.02, *η^2^_p_* = .46. Further analysis showed a main effect of Expertise for both AOI_NO_, *F*_(1,6)_ = 15.30, *p* = .008, and AOI_Outside_, *F*_(1,6)_ = 5.99, *p* < .05, indicating that for AOI_NO_, surgeons (*M* = 19.6%, *SD* = 3.5%) had higher percent dwell duration than novices (*M* = 11.6%, *SD* = 2.4%), Cohen’s *d* = 2.67, but for AOI_Outside_ surgeons (*M* = 59.6%, *SD* = 2.0%) had smaller percent dwell duration than novices (*M* = 72.1%, *SD* = 8.4%), Cohen’s *d* = −2.05.

When comparing percent dwell duration of novices at post-test **(Figure 4C, right)** and surgeons **(Figure 4D),** the Expertise by AOI ANOVA continued to show significant main effect of AOI, *F*_(2,12)_ = 152.20, *p* < .001, *η*^2^_p_ = .96, and a significant Expertise by AOI interaction, *F*_(2,12)_ = 8.05, *p* = .006, *η*^2^_p_ = .57. Further investigation demonstrated a significant main effect of Expertise on both AOI_TP_, *F*_(1,6)_ = 18.68, *p* = .005, and AOI_Outside_, *F*_(1,6)_ = 8.18, *p* = .03, with surgeons (*M* = 22.5%, *SD* = 4.9%) showing higher values than novices (*M* = 9.3%, *SD* = 4.2%), Cohen’s *d* = 3.46, for AOI_TP_, but surgeons (*M* = 59.6%, *SD* = 2.0%) showing lower values than novices (*M* = 74.4%, *SD* = 8.5%), Cohen’s *d* = −2.40 for AOI_Outside_. As such, the evidence indicates a decrease of gaze towards the target peg and an increase in gaze towards the next object that results from practice.

## Discussion

This study aimed to extend previous research applying eye-tracking technology to understand expertise and skill acquisition in laparoscopic surgery. Here, both behavioral and eye-tracking metrics were compared between experienced laparoscopic surgeons and novices, as novices underwent a multi-visit training protocol. The peg transfer task was selected to highlight bimanual coordination in laparoscopic surgery, while offering a validated approach to model individualized skill acquisition process through training. To adapt to eye-tracking constraints, the FLS peg transfer task was adjusted to fix the action sequence into a constant ordering, allowing objective AOIs to be defined. Results revealed that, although all the novices reached post-training behavioral performance that approximated their individualized learning plateaus and was statistically indistinguishable from that of the laparoscopic surgeons, their training experiences did not lead to expert gaze behaviors. In particular, whereas novices focused more on the AOI_Outside_, implying online visual sampling of object currently being moved, surgeons focused more on the AOI_TP_ and AOI_NO_, suggesting greater focus on feedforward sampling control. The evidence thereby supports the hypothesis that surgeons used a more proactive gaze strategies than novices, but not the hypothesis that training novices to expert-level behavioral performance would be accompanied by expert-like gaze pattern in a complex bimanual laparoscopic surgery task. The following discussion, therefore, addresses the implication of these findings, the strengths and weaknesses in this design and future directions for this research.

The current study utilized the validated FLS peg transfer task^3,15^ to characterize learning and explore group differences between novices and experienced surgeons. Each novice spent approximately three hours practicing the task, resulting in saturated behavioral learning according to both individual- and group-based results. As illustrated by the inverse model **(Figure 3),** on average novices were able to improve to 90% of their behavioral plateaus in 24 minutes, with 8% more improvement over the remaining training, leading to a learning rate ratio of 70 (i.e., 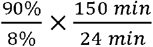). This decelerated learning rate across training time is unsurprising given the skill learning literature^20^, especially when the design is aimed to maximize the training volume (~3 hours) for possible alteration in corresponding gaze behaviors in task performance. However, the opportunity cost in offering this approximately saturated behavioral training volume becomes concerning when little change is observed in novices’ gaze patterns that consistently differ from that of surgeons in the laparoscopic simulation and in light of the restricted training hours available to surgical trainees^4^. The finding thus implies that surgical training programs focused on manual coordination alone cannot result in the complete development of eye-hand coordination patterns produced by surgeons.

One interesting observation from the current findings is that the surgeons showed highly consistent patterns of behavioral and eye-tracking results across individuals, as evidenced by relatively small standard deviations within the group. This illustrates a learned template with relatively greater feedforward visual sampling, where 40.4% of total dwell duration is distributed to AOIs that capture the upcoming targets in the action sequence for a given transfer. This finding is consistent with previous research and supports the view that visual expertise in surgery features flexibility in attentional distribution, accurate prediction of ensuing actions, and skillful use of parafoveal vision in controlling surgical tools^7,8,13^. Such a feedforward, externally-oriented, and autonomous attentional style has been shown to reduce electromyographical noise^21^, enhance short-loop reflexes in motor control^22,23^, and produce greater neural efficiency, which are earmarks of perceptual-motor expertise and may account for expert performance in surgery. Future research is thus encouraged to further explore the mechanisms underlying expert gaze pattern along these directions.

A close examination of the eye-tracking results indicates that the observed gaze pattern may bear different meanings when comparing surgeons to novices prior to and after training. Although novices always showed longer dwell duration in AOI_Outside_ than surgeons, their gaze in the AOI_NO_ and AOI_TP_ produced different profiles across the training. Specifically, relative to surgeons, novices demonstrated equivalent dwell duration on the target peg and less dwell duration on the next object at pre-test, whereas they showed shorter dwell duration on the target peg and equivalent dwell duration on the next object at post-test. Such an evidentiary pattern suggested a tradeoff in dwell duration between the two AOIs that were supposed to gauge feedforward visual sampling. One possible explanation from the novice’s standpoint is that, at pre-test, novices required greater monitoring of the dissectors and target objects in order to complete the transfer and successfully drop the object on the target peg. This challenge may have increased the proportion of gaze in AOI_TP_, which does not reflect feedforward visual sampling per se. As novices gained proficiency at the task, however, they may have been able to divert attention earlier from the AOI_TP_ to the AOI_NO_, reflecting an intention to work on the next object for faster task completion. The fact that AOI_NO_ did not differ between novices and surgeons after training may indicate the ability to shift, with practice, from sampling one step ahead on the target peg, to two steps ahead to view the next object in the sequence. A second possibility from the surgeon’s standpoint is that, because dropping the target object to the target peg in the task is designed to simulate the starting actions of suturing in laparoscopic surgery, surgeons may show the “cognitive slowing down”, reflecting a refocusing effort to increase attention towards a critical location in the surgical task due to professional experience^25,26^. Yet another possibility is that the truth lies in a combination of factors from both novices’ training and surgeons’ experience. The clarification of such subtle findings would merit future study.

The current findings have several implications for laparoscopic surgery research using eye-tracking. First, given the current evidence that novices do not completely develop the expert gaze pattern from manual practice alone, it may be advisable to also implement eye-tracking technology in surgical training programs so that the expertise gap can be closed not only on behavioral criteria but also on gaze pattern. Past studies have shown that this technique is able to differentiate experience surgeons from those of less experience when similar manual movement time was observed between the two cohorts^25^. Principles of such training design may include facilitating (a) awareness of the expert gaze pattern, (b) contrast between one’s own and expert gaze pattern, (c) recognition of targets in surgical actions, and (d) the self-regulatory “cognitive slow down”. Second, eye-tracking measures, particularly those on normalized scales such as the current percent dwell duration metric, seem more sensitive to surgical expertise than behavioral measures. Such an advantage of eye-tracking metrics has been suggested in other relevant research^8,13,27^, and can be explained by the high frequency (30 hz) of observation during data collection. Even though this large amount of eye-tracking data creates difficulties in management and processing, it may also increase the signal/noise ratio in detecting surgical expertise, suggesting the potential of eye-tracking metrics to be used in evaluating training program efficiency^28^. Finally, the current research made an effort to capture one’s visual attention in the peg transfer task by defining three AOIs based on analytical trials of object transfer. Although there might be more precise approaches, and particularly ones that precisely track gaze on the object currently in motion, the paradigm here was able to detect between-group differences providing a meaningful approach to studying learning and expertise as intended in this experiment.

## Conclusions

Novices achieved profound behavioral learning through training, leading to performance scores equivalent to experienced surgeons. Despite this, surgeons continued to demonstrate more feedforward visual sampling control by gazing at surgical targets of ensuing actions with such differences persisting even after training of novices. It can thus be inferred that, while traditional simulation-based laparoscopic skill training can improve novices’ behavioral task performance to a level similar to surgeons, differences in gaze pattern remain, motivating future surgical training programs to involve eye-tracking technology in its design and evaluation.

## Acknowledgments

Authors would like to thank Bob Wilson for help with the eye tracking technology and Dr. Ranjan Sudan for serving as study doctor for this study. The authors would also like to thank all of the participants for their time and effort and members of the Surgical Education and Activities Lab (SEAL) at Duke University, including Jennie Phillips and Layla Triplett, for their assistance with this research. The collaboration opportunity and funding for devices used in this research study was provided by Dr. Allan Kirk, Chair of the Duke University Department of Surgery, for which we would like to express our gratitude.

## Footnotes

1 The video can be found at https://www.youtube.com/watch?v=gAQPXHWqdXQ

2 Because of potential confounds introduced by certain AOIs locating in the pathways of task movement (see Figure 1B), a sensitivity analysis was performed using only Transfer 2 to 8 (i.e., removing Transfer 1 and Transfer 9 to 12). Results of this sensitivity analysis were consistent with those reported in the current Results section, supporting the robustness of current findings and indicating that the co-location did not affect the findings.

## Notes

**Funding:** This research was funded by grant support to L.G.A. through the United States Army Research Office [W911NF-15-1-0390].

**Conflict of interest:** All authors declare that they have no conflict of interest related to the research presented in this manuscript.

### Competing Interest Statement

The authors have declared no competing interest.

