## Supplemental Materials for "Skill acquisition and gaze behavior during laparoscopic surgical simulation"

**Supplement** **– Percent Dwell Count Results**

Novices Learning across Test Sessions

The AOI by Session ANOVA indicated a significant effect of AOI, *F*(1.67,6.67) = 19.45, *p* = .002, *η*^2^_p_ = .83, on the percent dwell count. Pairwise comparison revealed a significantly higher percent dwell count for AOI_Outside_ (*M* = 47.4%, *SD* = 6.4%) than for both AOI_NO_ (*M* = 31.0%, *SD* = 4.4%), *p* = .009, Cohen’s *d* = 1.75, and AOI_TP_ (*M* = 21.6%, *SD* = 5.2%), *p* < .001, Cohen’s *d* = 2.76. No significant findings on the main effect of Session or Session by AOI interaction were identified (*p*s > .15).

Surgeons versus Novices

The Expertise by AOI ANOVA comparing surgeons to novices at pre-test showed a significant effect of AOI, *F*(2,12) = 20.38, *p* < .001, *η*^2^_p_ = .77, and a significant Expertise $\times$ AOI interaction, *F*(2,12) = 6.37, *p* = .01, *η*^2^_p_ = .52 on the percent dwell count measure. Further analyses showed a significant simple main effect of Expertise for both AOI_NO_, *F*(1,6) = 10.67, *p* = .02, and AOI_Outside_, *F*(1,6) = 7.79, *p* = .03, indicating that for AOI_NO_ surgeons (*M* = 35.5%, *SD* = 2.9%) had higher percent dwell count than novices (*M* = 26.5%, *SD* = 4.2%), Cohen’s *d* = 2.52, but for AOI_Outside_ surgeons (*M* = 37.8%, *SD* = 1.7%) had lower percent dwell count than novices (*M* = 46.9%, *SD* = 5.3%), Cohen’s *d* = -2.31.

The Expertise by AOI ANOVA comparing surgeons to novices at post-test showed a significant effect of AOI, *F*(2,12) = 20.91, *p* < .001, *η*^2^_p_ = .78 on the percent dwell count measure, as well as a significant Expertise $\times$ AOI interaction, *F*(2,12) = 4.22, *p* =.04, *η*^2^_p_ = .41. Further analysis revealed a significant simple main effect of Expertise for AOI_TP_, *F*(1,6) = 8.43, *p* = .03, and a trending main effect of Expertise for AOI_Outside_, *F*(1,6) = 4.52, *p* = .078, showing that for AOI_TP_ surgeons (*M* = 26.7%, *SD* = 1.6%) had higher percent dwell count than novices (*M* = 18.2%, *SD* = 4.8%), Cohen’s *d* = 2.38, but for AOI_Outside_ surgeons (*M* = 37.8%, *SD* = 1.7%) had lower percent dwell count than novices (*M* = 47.4%, *SD* = 7.5%), Cohen’s *d* = -1.77.

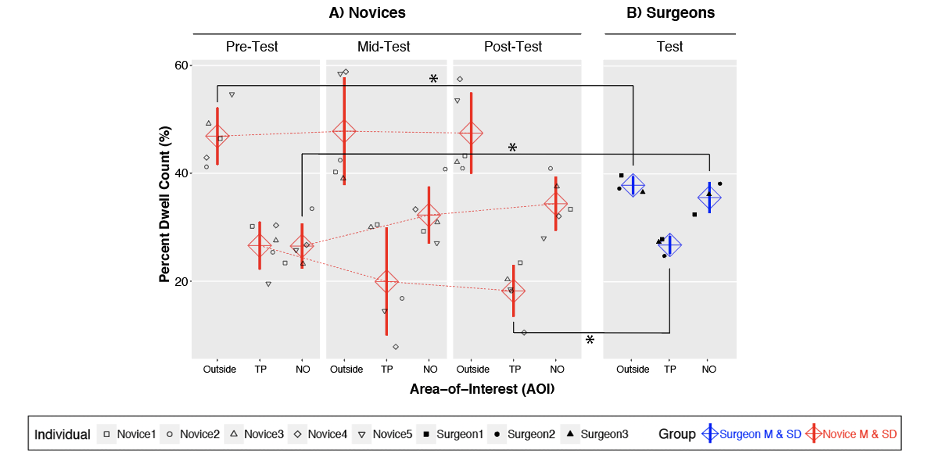

**Figure S1**. Percent dwell count (for both Individuals and groups) with important comparisons statistically marked. A) Novice percent dwell count for each AOI shown for each testing session. B) Surgeon percent dwell count for each AOI. **p* < .05.
